# More than just a ticket canceller: The mitochondrial processing peptidase matures complex precursor proteins at internal cleavage sites

**DOI:** 10.1101/2020.07.02.183996

**Authors:** Jana Friedl, Michael R. Knopp, Carina Groh, Eyal Paz, Sven B. Gould, Felix Boos, Johannes M. Herrmann

## Abstract

Most mitochondrial proteins are synthesized in the cytosol as precursors that carry N-terminal presequences. After import into mitochondria, these targeting signals are cleaved off by the mitochondrial processing peptidase MPP, giving rise to shorter mature proteins. Using the mitochondrial tandem protein Arg5,6 as a model substrate, we demonstrate that MPP has an additional role in preprotein maturation, beyond the removal of presequences. Arg5,6 is synthesized as a polyprotein precursor that is imported into the mitochondrial matrix and subsequently separated into two distinct enzymes that function in arginine biogenesis. This internal processing is performed by MPP, which cleaves the Arg5,6 precursor both at its N-terminus and at an internal site between the Arg5 and Arg6 parts. The peculiar organization and biogenesis of Arg5,6 is conserved across fungi and might preserve the mode of co-translational subunit association of the arginine biosynthesis complex of the polycistronic arginine operon in prokaryotic mitochondrial ancestors. Putative MPP cleavage sites are also present at the junctions in other mitochondrial fusion proteins from fungi, plants and animals. Our data suggest that, in addition to its role as “ticket canceller” for the removal of presequences, MPP exhibits a second, widely conserved activity as internal processing peptidase for complex mitochondrial precursor proteins.

## Introduction

All cellular processes are carried out by proteins, linear chains of amino acids that fold into three-dimensional structures. While the amino acid sequence of a protein is primarily determined by its DNA sequence, many proteins are additionally modified by proteolytic cleavage after their synthesis. Processing of polypeptides at their N-terminus is pervasive in both prokaryotic and eukaryotic proteomes. For instance, the amino-terminal methionine is removed from many polypeptides when they emerge from the ribosome and the new N-terminus is a crucial determinant of protein stability (Bradshaw et al., 1998, Varshavsky, 2011, Frottin et al., 2006). The majority of intracellular protein targeting signals are located at the N-terminus and cleaved off upon arrival at the correct cellular destination. For example, around two thirds of the nuclear-encoded mitochondrial proteins are synthesized as precursors that carry an N-terminal mitochondrial targeting sequence (MTS) or presequence which directs them to mitochondrial surface receptors (Bykov et al., 2020, Becker et al., 2019, von Heijne, 1986). These preproteins are imported into mitochondria via the translocases of the outer (TOM complex) and inner mitochondrial membrane (TIM23 complex) (Pfanner et al., 2019, Chacinska et al., 2009). Their MTS is cleaved by the mitochondrial processing peptidase (MPP). In some cases, the thereby generated new N-terminus of the polypeptide is further shortened by cleavage of single amino acids or short peptides by the proteases Icp55 or Oct1 before the matured protein folds into its native structure (Poveda-Huertes et al., 2017, Vögtle et al., 2011, Vögtle et al., 2009, Calvo et al., 2017, Naamati et al., 2009). Correct processing of precursor proteins in the matrix is crucial to maintain mitochondrial function and proteostasis. Dysfunctional preprotein maturation in mitochondria was observed in models of Alzheimer’s disease (Mossmann et al., 2014). It results in proteome instability, aggregation of incorrectly processed precursors and proteotoxic stress (Poveda-Huertes et al., 2020).

Internal cleavage of polypeptides is less frequent than processing from the N-terminus, but equally relevant for cellular physiology and organismal health. Examples of medically relevant polypeptides that undergo internal proteolytic processing during their biosynthesis include insulin or amyloid precursor protein (APP) (Steiner and Oyer, 1967, Müller et al., 2017). In most cases, peptides are removed to yield the mature form of a single protein. However, some genes encode fusion proteins that are synthesized as a single precursor and are then separated into distinct, functional proteins by proteolytic cleavage. A prominent example is ubiquitin, which is encoded as a fusion with subunits of the ribosome or as head-to-tail repeats of several ubiquitin monomers which are rapidly separated by deubiquitinating proteases (Finley et al., 1989, Ozkaynak et al., 1984, Gemayel et al., 2017). Polyproteins also frequently occur in viral genomes, including HIV and SARS-CoV-2, where the cleavage products often form protein complexes (Yost and Marcotrigiano, 2013, Zhang et al., 2020, Krichel et al., 2020). In eukaryotic genomes, such an organization is rare, despite its obvious advantage of stoichiometric co-expression of functionally related proteins.

Here we report about a notable exception: The *ARG5,6* gene of *Saccharomyces cerevisiae* encodes both the acetylglutamate kinase (Arg6) and the acetylglutamyl-phosphate reductase (Arg5), two enzymes that catalyze the second and third step of arginine biosynthesis in the mitochondrial matrix (Minet et al., 1979). These two enzymes are synthesized as a single precursor protein that is post-translationally cleaved into separate polypeptides. Arg6 and Arg5 then form a complex together with the acetylglutamate synthase Arg2 (Pauwels et al., 2003, Abadjieva et al., 2001). Here, we investigate the biogenesis of Arg5,6 in more detail and identify MPP as the protease that is responsible for processing of the precursor into two functional enzymes. We demonstrate that Arg6 and Arg5 can be imported into mitochondria separately, where they still form a functional enzyme. However, its organization as a composite precursor that is matured by MPP is highly conserved across fungi. These findings broaden our view on MPP as processing peptidase of more general role in mitochondrial preprotein maturation, reaching beyond its canonical function of removing mitochondrial targeting signals.

## Results

### Arg5,6 is processed by MPP in the mitochondrial matrix

The acetylglutamate kinase (Arg6) and acetylglutamyl-phosphate reductase (Arg5) are located in the mitochondrial matrix, where they catalyze the second and third step of the biosynthesis of arginine from glutamate (Fig. 1A) (Minet et al., 1979, Morgenstern et al., 2017). The composite precursor protein that is synthesized from the *ARG5,6* gene contains a mitochondrial targeting sequence (MTS) (Vögtle et al., 2009), which was suggested to direct the single precursor into mitochondria where it is subsequently cleaved into two separate polypeptides (Boonchird et al., 1991b). Indeed, when we expressed Arg5,6 with a C-terminal hemagglutinin (HA) tag, immunoblotting of cell lysates against the HA epitope revealed only a single band at around 40 kDa, much less than the expected mass of 90 kDa of the composite precursor protein (Fig. 1B). Obviously, the precursor is rapidly and efficiently cleaved *in vivo*, which yields a C-terminal Arg5 fragment.

**Figure 1.**
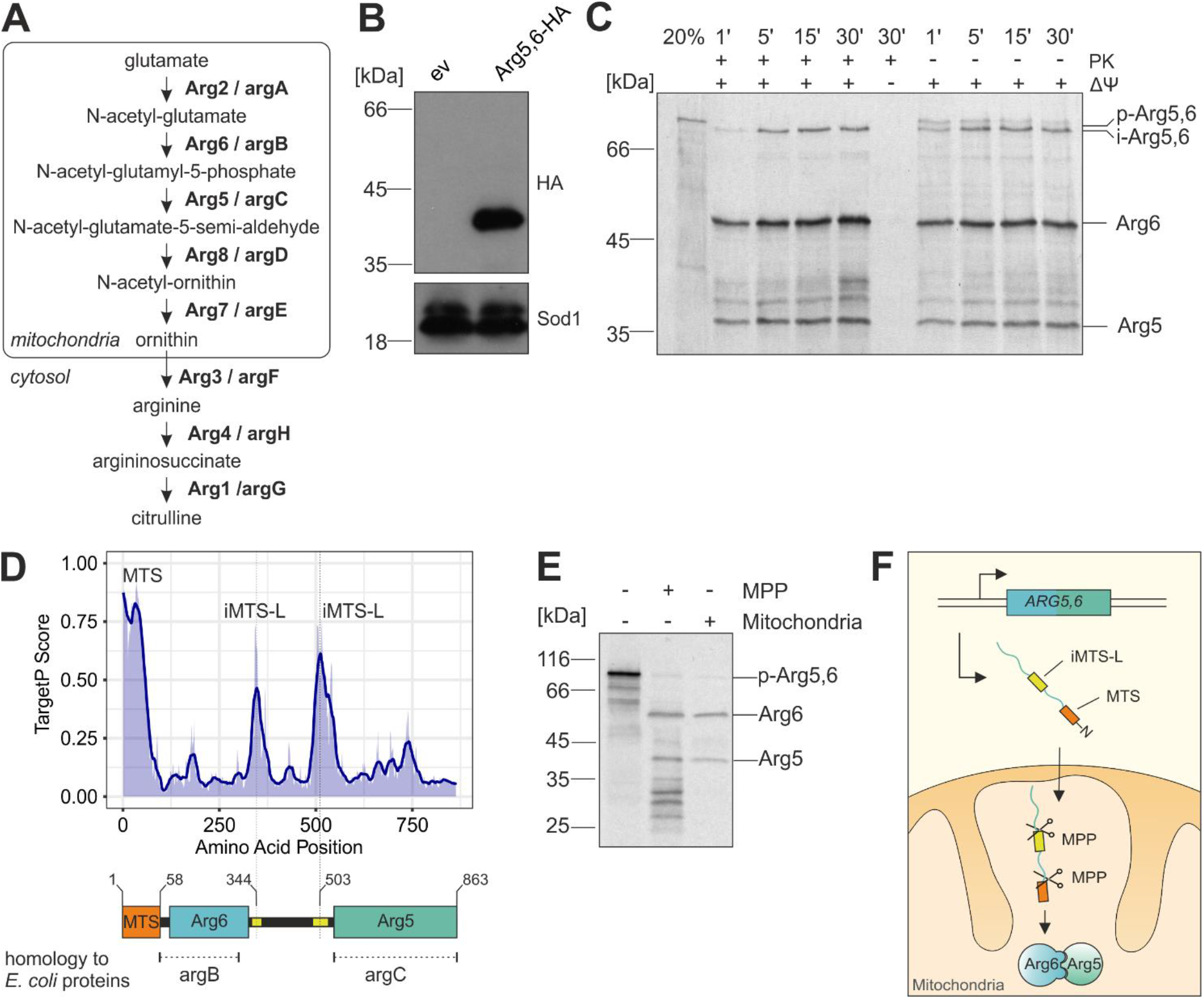
Arg5,6 is a composite mitochondrial precursor that is processed twice by MPP in the mitochondrial matrix. **A,** Schematic representation of arginine biosynthesis in *S. cerevisiae*. Shown in bold are the enzymes that catalyze the respective step as well as their *E. coli* orthologs. **B,** When Arg5,6 was C-terminally HA-tagged, immunoblotting revealed a single band at 40 kDa, indicating proteolytic cleavage of the 90 kDa precursor protein. **C,** Radiolabeled Arg5,6 precursor was incubated with isolated mitochondria for the indicated times and analyzed by SDS-PAGE and autoradiography. Non-imported material was digested with proteinase K (left half). 20% of the total lysate used per import lane was loaded for control. The membrane potential (Δψ) was dissipated with VAO (valinomycin, antimycin, oligomycin). p, precursor, i, intermediate. **D,** Arg5,6 was subjected to TargetP profiling. High values indicate regions within the protein which structurally resemble mitochondrial presequences. **E,** His-tagged MPP was expressed and purified from *E. coli*. Radiolabeled Arg5,6 precursor was incubated with isolated mitochondria for 15 min or purified MPP for 90 min. The processing of Arg5,6 was analyzed by SDS-PAGE, Western blotting and autoradiography. **F,** Arg5,6 is imported into the mitochondrial matrix and cleaved twice by MPP: once at the N-terminus to remove the presequence, and once internally at an iMTS-L to separate Arg5 and Arg6.

In order to analyze the mechanistic basis of this unusual biogenesis, we sought to reconstitute the biogenesis and processing of Arg5,6 in an *in vitro* system. To this end, we synthesized radiolabeled Arg5,6 precursor in reticulocyte lysate and incubated it with isolated yeast mitochondria. We observed that the precursor of around 90 kDa was efficiently processed to a slightly smaller intermediate form – indicating the removal of the N-terminal MTS – and further into several smaller fragments, most prominently two polypeptides of around 40 and 50 kDa, which correspond to the C-terminal Arg5 and the N-terminal Arg6, respectively (Fig. 1C). The intermediate as well as the mature polypeptides, but not the Arg5,6 precursor were protected from digestion with externally added proteinase K. This shows that these polypeptides were translocated across the mitochondrial outer membrane. Both import and processing were dependent on the mitochondrial inner membrane potential (Δψ) as expected for an MTS-containing matrix protein (Sato et al., 2019, Schendzielorz et al., 2017, Garg and Gould, 2016). We conclude that Arg5,6 is imported into the mitochondrial matrix via the presequence pathway and cleaved into separate polypeptides inside mitochondria.

How is the Arg5,6 precursor processed to give rise to the Arg6 and Arg5 enzymes? A number of proteases in the mitochondrial matrix are described (Quiros et al., 2015, Veling et al., 2017). However, most of them are either implicated in degradation and turnover of proteins (such as the Lon protease Pim1) or known to remove short peptides or single amino acids from the N-terminus of mitochondrial precursor proteins (such as Oct1 or Icp55), but not for internal cleavage of proteins into two mature parts (Vögtle et al., 2009, Poveda-Huertes et al., 2017, Vögtle et al., 2011, Wagner et al., 1994, Suzuki et al., 1994). For MPP, the peptidase responsible for the cleavage of the N-terminal MTS, a notable exception was recently reported: The composite precursor protein Atp25 contains internal MPP cleavage sites at which the protein is split into two functionally unrelated polypeptides (Woellhaf et al., 2016). In Atp25, the internal MPP cleavage sites coincide with a sequence stretch that structurally mimics the properties of an N-terminal MTS. We asked whether MPP might also be involved in the processing of Arg5,6 and subjected its amino acid sequence to an *in silico* prediction of such internal MTS-like sequences (iMTS-Ls) using an adapted version of the TargetP algorithm (Boos et al., 2018, Emanuelsson et al., 2007). In fact, in addition to the N-terminal MTS, two internal regions with high TargetP score were detected around amino acid positions 344 and 503 (Fig. 1D). The latter one would fit to the molecular masses of the Arg6 and Arg5 proteins observed in our *in vitro* system. To directly test whether MPP can cleave the Arg5,6 precursor, we purified MPP from *E. coli* expressing His-tagged Mas1 and Mas2 (the two subunits of MPP). Incubation of radiolabeled Arg5,6 precursor protein with MPP resulted in the formation of smaller fragments whose size perfectly matched those that were generated after import into isolated mitochondria (Fig. 1E). Proper processing was blocked when EDTA was added to the reaction, which inhibits the metalloprotease MPP by chelating divalent cations (Suppl. Fig. 1A) (Luciano et al., 1998).

We conclude that Arg5,6 is imported into the mitochondrial matrix and processed twice by MPP. A first cleavage removes the N-terminal MTS and a second cleavage at an iMTS-L separates the Arg6 and Arg5 enzymes (Fig. 1F).

### Arg5 and Arg6 can be imported into mitochondria separately and complement the *arg5,6* deletion mutant

The unusual biogenesis of Arg5,6 prompted us to ask whether Arg6 and Arg5 can also be imported separately. Therefore, we created truncated versions of the *ARG5,6* gene which contain only the N-terminal Arg6 with its MTS (Arg6^1-502^) or only the C-terminal Arg5, starting at the first (Arg5^344-863^) or the second iMTS-L (Arg5^503-863^). For the latter two, we also generated variants which additionally carry the well-characterized presequence of ATP synthase subunit 9 from *Neurospora crassa* (Su9-Arg5^344-863^ and Su9-Arg5^503-863^) (Fig. 2A).

**Figure 2.**
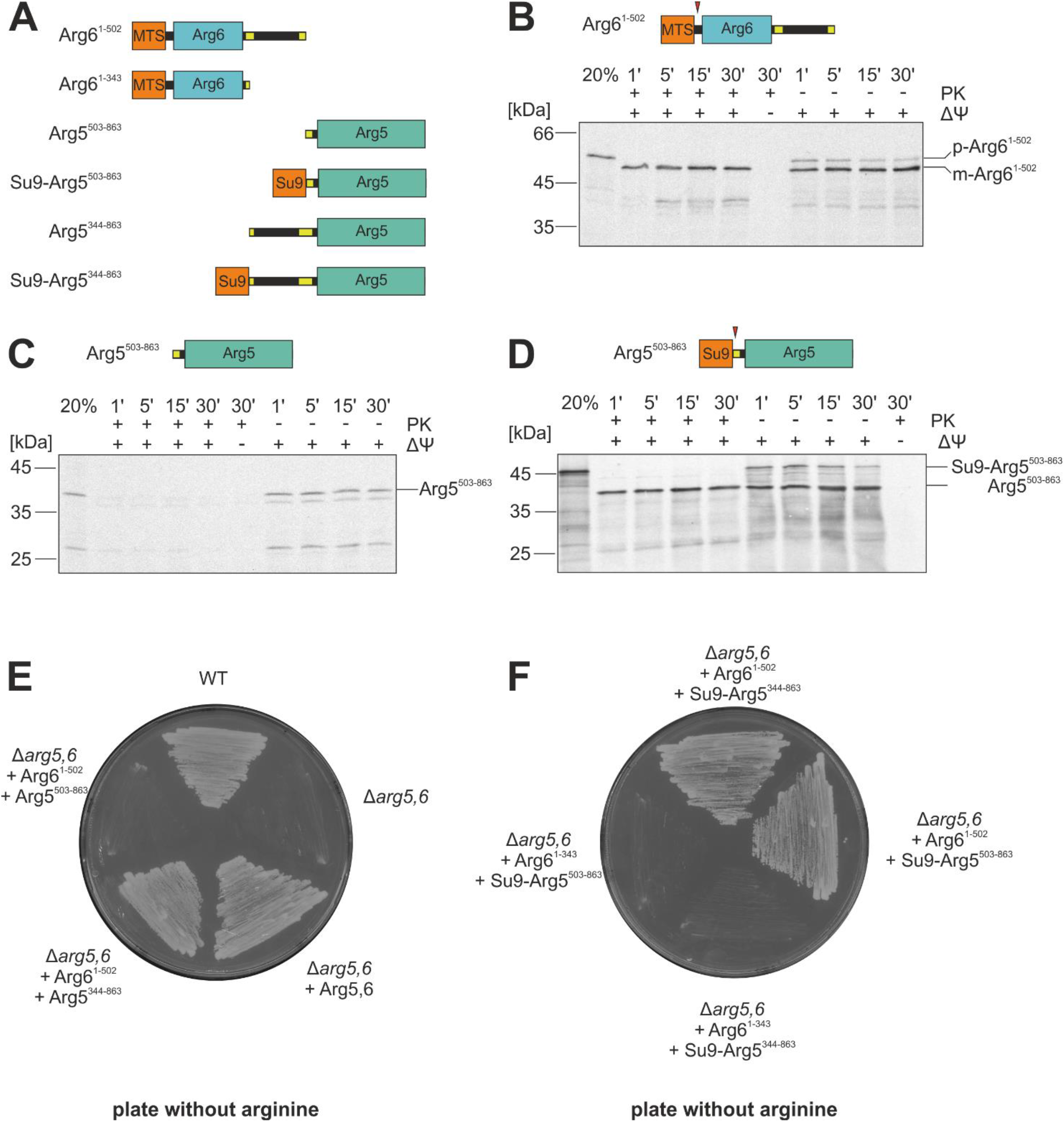
Arg5 and Arg6 can be imported separately *in vitro* and *in vivo*. **A,** Overview of truncated Arg5,6 variants used in this study. Su9, presequence of *N. crassa* subunit 9. **B-D,** Radiolabeled precursor proteins of Arg6^1-502^, Arg5^503-862^ and Su9-Arg5^503-862^ were incubated with isolated mitochondria for the indicated times and analyzed by SDS-PAGE and autoradiography. Non-imported material was digested with proteinase K (left half). 20% of the total lysate used per import lane is loaded for control. The membrane potential (Δψ) was depleted with VAO. Red arrowheads indicate processing sites. p, precursor, m, mature. **E-F,** Yeast cells that lack endogenous Arg5,6 (Δ*arg5,6*) were transformed with plasmids for expression of the indicated Arg5,6 variants and streaked out on plates containing minimal growth medium without arginine.

Radiolabeled proteins were synthesized *in vitro* and incubated with isolated mitochondria to test their import competence. As expected, Arg6^1-502^ was efficiently imported and its MTS was cleaved (Fig. 2B). The shorter Arg5 variant (Arg5^503-863^) did not reach a protease-protected compartment and, thus, was not imported into mitochondria (Fig. 2C). However, N-terminal fusion of the Su9 presequence completely restored import of Arg5^503-863^ (Fig. 2D). Hence, import of Arg5 and Arg6 into mitochondria is in principle possible also for separated polypeptides, at least in the *in vitro* assay used here.

We next tested whether Arg6 and Arg5 can be imported separately *in vivo* and function in arginine biosynthesis. The deletion of the *ARG5,6* gene renders yeast cells auxotrophic for arginine. We expressed either the full length Arg5,6 precursor or combinations of separate Arg6 and Arg5 variants in a Δ*arg5,6* deletion mutant. If Arg6 and Arg5 make their way into mitochondria and acquire a functional conformation, arginine prototrophy should be restored. When we streaked out these cells on plates with minimal growth medium lacking arginine, we observed growth for the wildtype and the Δ*arg5,6* mutant complemented with full length Arg5,6, but not for Δ*arg5,6* carrying only an empty plasmid, as expected (Fig. 2E).

The mutant expressing both Arg6^1-502^ and the shorter Arg5^503-863^ variant was not able to grow without arginine (Fig. 2E). However, when the presequence of Su9 was fused to the short Arg5^503-863^, cells regained arginine prototrophy (Fig. 2F). All strains grew on plates containing arginine, showing that the Arg5^503-863^ protein has no toxic gain-of-function effect when residing in the cytosol (Suppl. Fig. 1B,C). Growing the strains in liquid medium lacking arginine confirmed the results obtained on plates and additionally demonstrated that the growth rate of the strain expressing Arg6^1-502^ and Su9-Arg5^503-863^ is comparable to that of the wildtype (Suppl. Fig. 1D), at least under the overexpression conditions used here. This indicates that separate expression of Arg6 and Arg5 is, in principle, possible without adverse effects on cellular fitness.

A truncated version of Arg6 (Arg6^1-343^) did not complement the deletion mutant with any of the Arg5 variants, even though this variant contains the entire region of homology to the bacterial acetylglutamate kinase argB from *E. coli* (Fig. 2F). Apparently, the amino acids 344 to 502 are functionally relevant for enzyme activity and not merely a spacer or linker between Arg5 and Arg6.

Taken together, both Arg6 and Arg5 can be imported separately into mitochondria *in vitro* and *in vivo* and are functional in arginine biosynthesis, as long as mitochondrial localization is conferred by appropriate N-terminal targeting signals.

### Internal processing of Arg5,6 by MPP requires an N-terminal presequence

Surprisingly, cells expressing Arg6^1-502^ and the longer Arg5^344-863^ could grow without arginine, indicating that this longer Arg5 variant can be imported even without fusion of an additional presequence (Fig. 2E). Indeed, Arg5^344-863^ was imported into isolated mitochondria, albeit with low efficiency (Fig. 3A). Addition of proteinase K to this import reaction resulted in the appearance of lower-running bands that were absent without protease, indicating that a portion of the precursor translocated only partially across the outer membrane. This is in agreement with earlier observations that iMTS-Ls can confer targeting to mitochondria when presented at the N-terminus, but are not necessarily able to drive complete translocation (Backes et al., 2018, Baker and Schatz, 1987). As expected, fusion of the Su9 presequence to Arg5^344-863^ resulted in more efficient translocation (Fig. 3B).

**Figure 3.**
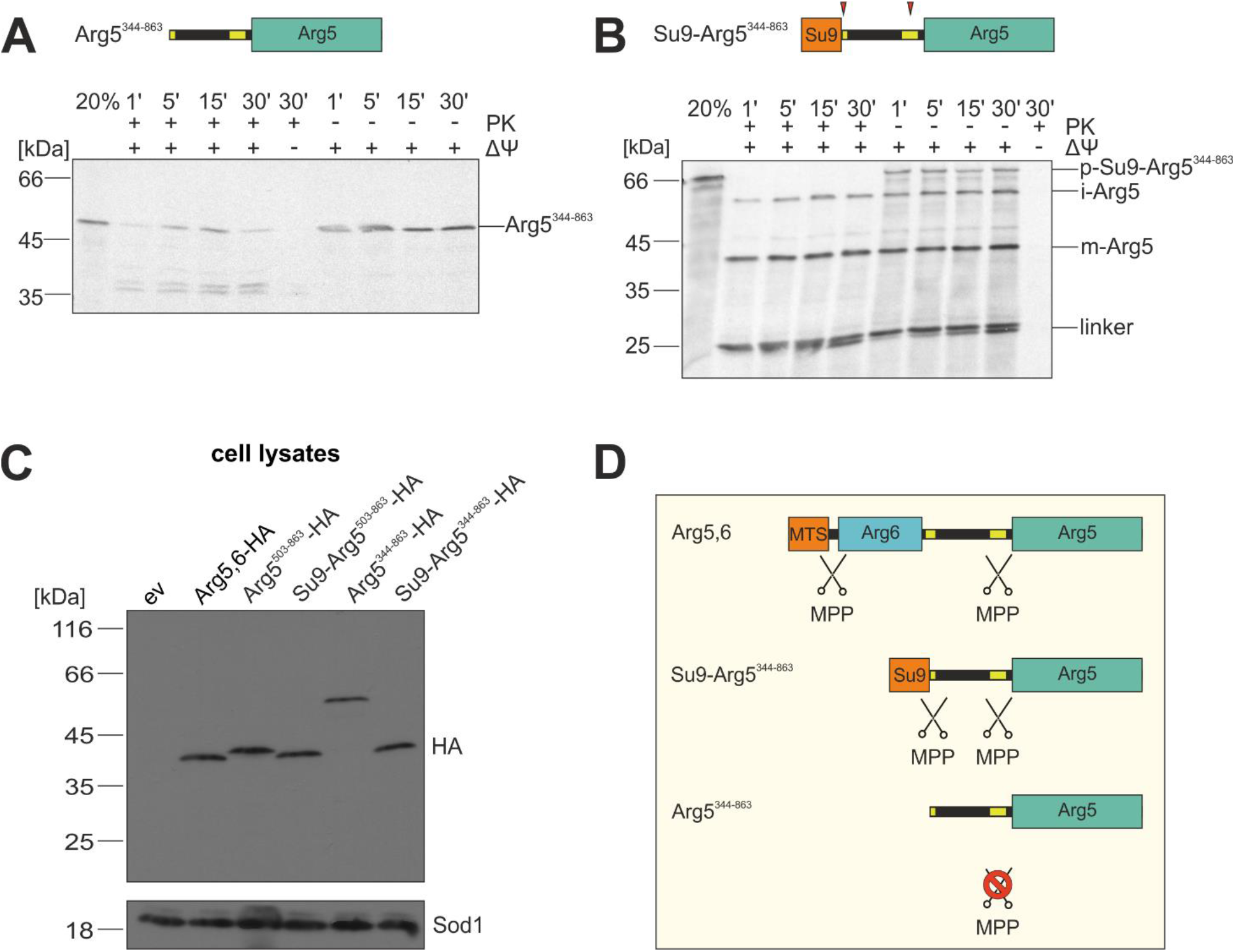
MPP requires a strong N-terminal MTS for internal processing of precursor proteins. **A-B,** Radiolabeled precursor proteins of Arg5^344-862^ and Su9-Arg5^344-862^ were incubated with isolated mitochondria for the indicated times and analyzed by SDS-PAGE and autoradiography. Non-imported material is digested with proteinase K (left half). 20% of the total lysate used per import lane is loaded for control. The membrane potential (Δψ) was depleted with VAO. Red arrowheads indicate processing sites. p, precursor, i, intermediate, m, mature. **C,** Yeast cells expressing indicated variants of Arg5,6, all carrying a C-terminal HA tag, were lysed and protein extracts were analyzed by SDS-PAGE and immunoblotting directed against the HA epitope or Sod1 as a loading control. ev, empty vector. **D,** MPP cleaves the Arg5,6 precursors at its internal processing site only if they carry a *bona fide* N-terminal presequence.

Interestingly, even the fully imported Arg5^344-863^ was not processed inside mitochondria, while for Su9-Arg5^344-863^, prominent lower-running bands were observed. Their sizes fit those expected for the intermediate form after removal of the Su9 presequence (around 60 kDa), the completely matured Arg5 after cleavage at the second MPP site (around 40 kDa) and the cleaved “linker” between the Su9 presequence and Arg5 (around 25 kDa). This suggests that an internal MPP cleavage at an iMTS-L requires a *bona fide* N-terminal MTS.

To test whether this holds true also *in vivo*, we expressed HA-tagged Arg5,6 as well as Arg5 variants in yeast cells and analyzed cell lysates by immunoblotting against the HA epitope. In agreement with the *in vitro* import experiments, both Su9-Arg5^344-863^ and Su9-Arg5^503-863^ were processed and yielded a band at the same molecular weight as full length Arg5,6. In contrast, Arg5^344-863^ and Arg5^503-863^ were exclusively present in their unprocessed forms (Fig. 3C). Hence, even though Arg5^344-863^ is imported into mitochondria, its iMTS-L is not recognized by MPP. Obviously, internal MPP cleavage sites are dependent not only on the presence of a particular motif and its immediate context within the amino acid sequence, but also on more distant features of the polypeptide, such as the N-terminal presequence (Fig. 3D).

### The *ARG5,6* genes are fused in fungi, separated in algae, and encoded polycistronically in gamma-proteobacteria

Tandem organization of functionally related genes is rare in eukaryotes. In contrast, genomic co-localization of functionally related genes is pervasive in prokaryotes. The *E. coli* proteins argB and argC are homologous to Arg6 and Arg5, respectively, and are organized in an operon, i.e. they are transcribed polycistronically (Piette et al., 1982). We used the sequences of argB and argC of *E. coli K-12* as query sequences to search a prokaryotic database of 5,655 organisms (Suppl. Table 1A). 3,666 species were identified that encode exactly one copy each of argB and argC (Suppl. Table 1B), mainly distributed among Proteobacteria, Firmicutes, Actinobacteria and Cyanobacteria. Among the 3,666 prokaryotes, 882 strains (848 of which were gamma-proteobacteria) encode the two open reading frames in immediate proximity, and hence presumably express them on a polycistronic mRNA, (Suppl. Table 1C).

Even though we cannot directly trace back the evolutionary history of Arg5,6 of *Saccharomyces cerevisiae*, these findings support the idea that this eukaryotic two-gene cluster evolved from syntenic genes in the eukaryotic ancestor. Nevertheless, Arg5,6 represents an exceptional case. Only very few proteins with a similar fusion structure are known in baker’s yeast, and for none of them do the separate proteins function in the same pathway (Woellhaf et al., 2016, Finley et al., 1989, Ozkaynak et al., 1987). However, the Arg5,6 homolog *arg-6* in *Neurospora crassa* also encodes both acetylglutamate kinase and acetylglutamyl-phosphate reductase which are separated by proteolytic cleavage inside mitochondria (Parra-Gessert et al., 1998, Gessert et al., 1994). We therefore asked whether the composite structure of Arg5,6 is generally conserved across eukaryote species. To this end, we searched a dataset of 150 eukaryotes for homologs of the yeast composite Arg5,6 precursor and of its post-translationally processed proteins Arg5 and Arg6 (Suppl. Table 1D). While 82 genomes returned no hits, we identified specific patterns within the diamond BLASTp hits of 36 eukaryotes (Fig. 4A, Suppl. Fig. 2, Suppl. Table 1E). In 28 species, all three query sequences hit the same subject, indicating the presence of an *ARG5,6* gene fusion homolog. With the sole exception of *Phytophthora sojae*, the fusion homologs were exclusively found among fungi. In eight species, Arg5 and Arg6 each hit a single independent sequence, while Arg5,6 hit both of those sequences, which indicates that separate genes encode for Arg5 and Arg6. Again with a single exception (*Fusarium graminearum* PH-1), all of these organisms were algae, comprising both green and red algae. In 32 species we identified mixed blast patterns (Suppl. Table 1F) that were not further investigated. We also predicted the intracellular localization of all proteins via TargetP. Strikingly, mitochondrial localization was assigned to almost all fusion proteins of fungi, whereas most separate proteins in algae were predicted to be imported into chloroplasts (Fig. 4B).

**Figure 4.**
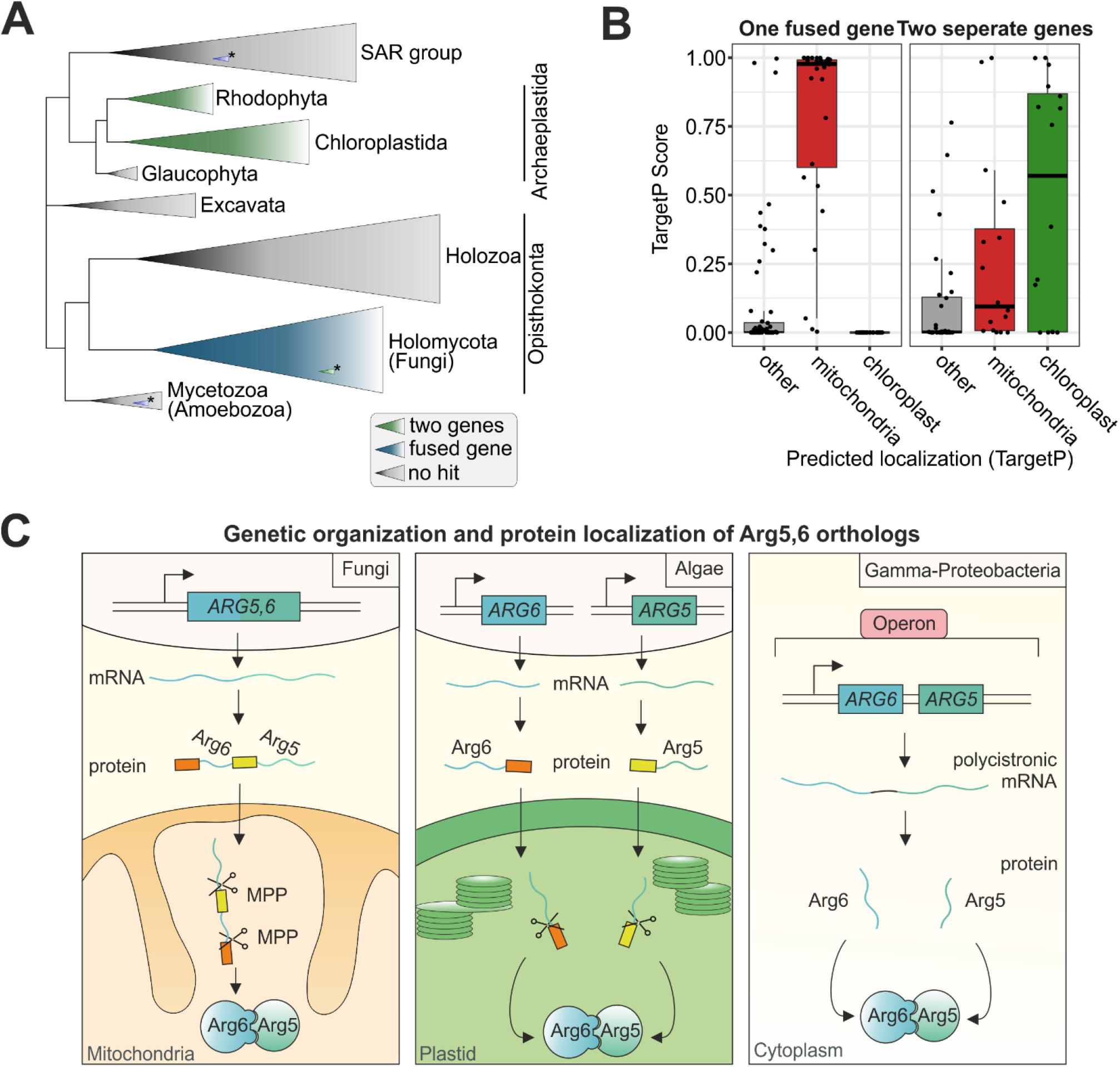
Tandem organization and mitochondrial localization of Arg5,6 is conserved in fungi, whereas algae synthesize two separate proteins that localize to their plastids. **A,** A database of 150 eukaryotes was searched for homologs of full length Arg5,6 and the separate Arg5 and Arg6 proteins of *S. cerevisiae*. Clades in which Arg5,6 homologs were encoded as a single fusion protein are coloured in blue, clades in which two separate Arg6 and Arg5 genes were found are coloured in green. Grey, no Arg5,6 homologs were found in these clades. **B,** In species that encode Arg5,6 as fusion protein, TargetP predicts the protein to be localized to mitochondria. In species with separate genes for Arg5 and Arg6, localization is predicted to be plastidal. **C,** In fungi, Arg5,6 is a fusion protein which is matured into two separate enzymes by MPP in the mitochondrial matrix. In contrast, algae encode two separate proteins that localize to their plastids. Gamma-proteobacteria encode Arg5 and Arg6 polycistronically.

In summary, in fungi the acetylglutamate kinase and acetylglutamyl-phosphate reductase are generally encoded as a fusion protein, which is imported into mitochondria and processed twice by MPP to remove its presequence and gives rise to two functional enzymes. In contrast, algae encode two separate proteins which are individually imported into chloroplasts. Gamma-proteobacteria express the genes from one polycistronic RNA (Fig. 4C).

### Internal precursor processing by MPP is conserved among eukaryotes

Since mitochondrial localization correlates well with the tandem structure of Arg5,6 homologs, we wondered whether the intramitochondrial processing into two separate enzymes by MPP might also be conserved in these species. To address this, we calculated the iMTS-L profiles of all fusion homologs of Arg5,6. In fact, we observed a common pattern across nearly all species: Besides the N-terminal MTS, our algorithm detected one conserved iMTS-L around amino acid position 500 (Fig. 5A). Hence, all these fusion proteins harbor a potential MPP cleavage site precisely at the junction between the Arg5 and Arg6 parts.

**Figure 5.**
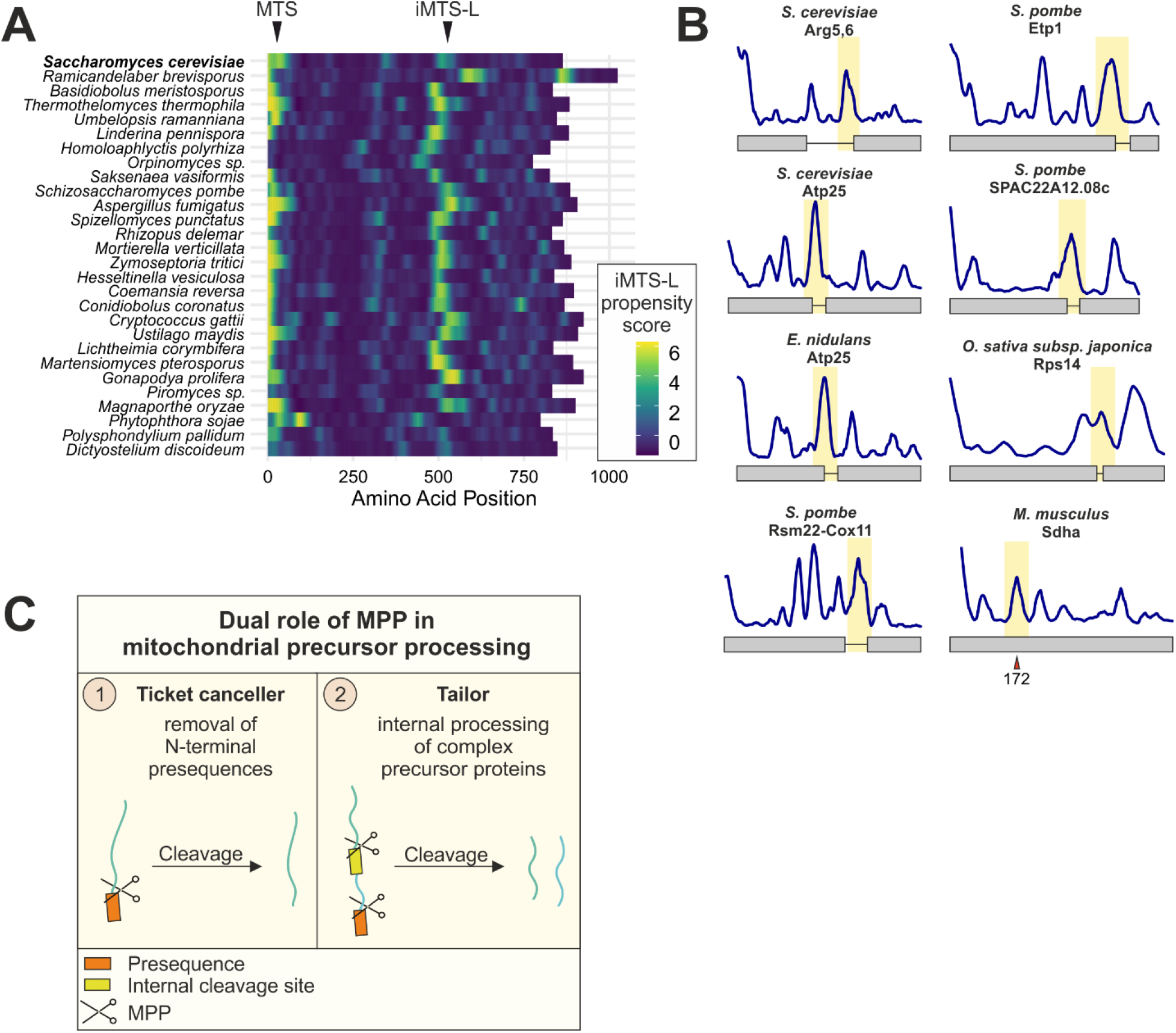
MPP functions not only as presequence peptidase, but also has a conserved processing activity at internal cleavage sites of composite precursor proteins. **A,** The iMTS-L at which MPP cleaves the Arg5,6 precursor in *S. cerevisiae* is conserved among species that encode Arg5,6 as fusion protein. Shown are iMTS-L propensity profiles along the sequence of Arg5,6 fusion protein homologs. **B,** Previously described mitochondrial polyproteins from different species harbor an iMTS-L at the position of the junction between the fused polypeptides, indicating potential MPP cleavage sites. Sdha is no polyprotein, but an alternative N-terminus (red arrowhead) was recently identified, presumably the result of a posttranslational cleavage (Calvo et al., 2017). **C,** In addition to its canonical role in presequence removal from mitochondrial precursor proteins, MPP has a second, conserved function. It recognizes internal cleavage sites and processes complex precursor proteins into separate polypeptides.

This remarkable conservation of protein structure and processing inspired us to ask whether MPP could be responsible for internal cleavage of other mitochondrial proteins. Apart from Arg5,6 and its homologs, some other mitochondrial proteins with composite structure have been described in different species. These include proteins from different fungi (Atp25 from *S. cerevisiae* and *E. nidulans* and Etp1, Rsm22-Cox11 and the uncharacterized SPAC22A12.08c from *S. pombe*) and plants (RPS14 from *O. sativa subsp. japonica*) (Oshima et al., 2005, Woellhaf et al., 2016, Khalimonchuk et al., 2006). We subjected these mitochondrial fusion proteins to our iMTS-L profiling analysis and indeed found prominent iMTS-Ls at each of their junction sites (Fig. 5B).

## Discussion

The organization as fusion protein is an elegant solution to confer mitochondrial targeting of two enzymes that reside in the same compartment and even act in subsequent steps of a biochemical pathway. It is still remarkable that this organization of Arg5,6 was retained during evolution even in distantly related organisms, indicating that there exists a strong constraint that maintained this organization for more than a billion years of evolution. To our knowledge, this is the only example of a fusion of two functionally related proteins whose organization is so widely conserved across eukaryote species. Eukaryotic genomes typically strongly disfavor even “milder” variants of physical coupling of genes, such as an operon-like organization which is pervasively present in prokaryotes, but basically absent in most eukaryotes. Interestingly, also events of horizontal gene transfer from bacteria to eukaryotes were accompanied by progressive loss of the polycistronic organization of the genes (Kominek et al., 2019). What might be the reason that the peculiar tandem structure of Arg5,6 survived several million years of evolution?

Since Arg6 and Arg5 form a complex, it is conceivable that the tandem structure of the precursor might facilitate their assembly. Many cytosolic protein complexes assemble cotranslationally, and this early-onset interaction between the partner subunits is crucial for function and, more generally, for maintenance of proteostasis (Shiber et al., 2018, Schwarz and Beck, 2019). Cotranslational assembly of nuclear encoded mitochondrial proteins is hindered by the additional translocation step across two mitochondrial membranes. Therefore, coupling the assembly with the import of proteins into the mitochondrial matrix might represent the closest approximation to cotranslational assembly that is physically possible.

Besides its canonical role as “ticket canceller” that clips targeting signals, the mitochondrial processing peptidase MPP obviously also possesses a “tailor” activity for internal processing of several precursor proteins (Fig. 5C). This property is conserved across the fungi, plant and potentially also animal kingdoms. Internal MPP cleavage requires a proximal recognition motif which appears to be an iMTS-L, but remarkably also a strong N-terminal presequence. This suggests that in order to access internal cleavage sites, MPP has to be loaded onto a precursor as soon as it emerges from the TIM23 channel. MPP might then “scan” the still unfolded polypeptide for cleavage sites. A protein with a strong presequence will efficiently recruit MPP, which enables subsequent internal cleavage at an iMTS-L before this is buried by protein folding. In analyses of the N-proteome from yeast, mouse and human mitochondria, a surprising variability in the N-termini of many proteins were observed, sometimes more than 100 amino acids downstream of the annotated start (Calvo et al., 2017, Vögtle et al., 2009, Vaca Jacome et al., 2015). Some of these isoforms might be generated by “leaky” internal MPP cleavage. For instance, the mouse protein Sdha has an alternative N-terminus at position 172 which coincides with an iMTS-L (Fig. 5B). It will be exciting to elucidate the biogenesis and physiological role of such protein isoforms in future research. Its unusual biogenesis might render Arg5,6 a valuable model substrate to explore the mechanisms that confer specificity of internal MPP cleavage.

## Materials and Methods

### Yeast strains and plasmids

All yeast strains used in this study were based on the WT strain BY4742 (Winston et al., 1995). Unless indicated differently, strains were grown on synthetic medium (0.17% yeast nitrogen base and 0.5% (NH4)2SO4) containing 2% glucose.

The Arg5,6-coding region or a fragment of it was amplified by PCR and cloned into pGEM4 (Promega, Madison, WI) using the EcoRI and BamHI restriction sites. For construction of the Arg5,6-HA, Arg5^344-863^-HA and Arg5^503-863^-HA versions, the corresponding sequence without stop codon was cloned into the expression plasmid pYX142, which harbors a constitutive *TPI* promoter upstream and the sequence of a hemagglutinin (HA) tag downstream of the multiple cloning site. For expression of Arg6^1-343^-HA and Arg6^1-502^-HA, pYX122 vectors were used which differ from pYX142 in the selectable marker. The sequence of the Su9 presequence was amplified from a Su9-DHFR plasmid and cloned into the EcoRI site of the pYX142 plasmid carrying the *ARG5* inserts to yield the Su9-Arg5^344-863^-HA and Su9-Arg5^503-863^-HA variants.

### Isolation of yeast mitochondria

Isolation of mitochondria was performed essentially as described (Saladi et al., 2020). Yeast cells were grown in YPGal medium (1% yeast extract, 2% peptone, 2% galactose) to an OD600 of 0.7-1.3, harvested, washed with water and resuspended in MP1 buffer (10 mM DTT, 100 mM Tris). Cells were incubated for 10 min at 30°C, pelleted (5 min at 4,000 x g at RT) and washed with 1.2 M sorbitol. The cell was digested in MP2 buffer (1.2M sorbitol, 20mM KH2PO4 pH 7.4) supplemented with 3mg zymolyase/g wet weight (Seikagaku Corporation) for 1 h at 30°C. The spheroplasts were harvested, resuspended in ice-cold homogenization buffer (0.6 M sorbitol, 1 mM EDTA pH 8, 1 mM phenylmethanesulfonyl fluoride (PMSF), 10 mM Tris-HCl pH 7.4, 0.2% fatty acid-free BSA) and lysed by douncing 10 times in a cooled potter homogenizer.

The homogenate was centrifuged for 5 min at 3,500 x g at 4°C to separate cell debris and nuclei from organelles. The mitochondrial fraction was isolated by centrifugation of the supernatant from the previous step for 12 min at 12,000 x g at 4°C. The crude mitochondrial pellet was gently resuspended in SH buffer (0.6M sorbitol, 20mM HEPES/KOH pH 7.4), centrifuged for 5 min at 4,000 x g at 4°C, recovered from the supernatant by centrifugation for 12 min at 12,000 x g at 4°C and finally resuspended in SH buffer. The protein concentration of the mitochondrial suspension was determined by a Bradford assay and mitochondria were diluted to a final concentration of 10 mg/ml protein with ice-cold SH buffer, aliquoted, frozen in liquid nitrogen and stored at −80°C.

### Import of radiolabeled proteins into isolated mitochondria

Import reactions were essentially performed as described previously (Peleh et al., 2015) in the following import buffer: 500 mM sorbitol, 50 mM Hepes, pH 7.4, 80 mM KCl, 10 mM magnesium acetate, and 2 mM KH2PO4. Mitochondria were energized by addition of 2 mM ATP and 2 mM NADH before radiolabeled precursor proteins were added. To dissipate the membrane potential, a mixture of 1 μg/ml valinomycin, 8.8 μg/ml antimycin, and 17 μg/ml oligomycin was added to the mitochondria. Precursor proteins were incubated with mitochondria for different times at 25°C before nonimported protein was degraded by addition of 100 μg/ml proteinase K.

### Growth Assays

Growth curves were performed automated in a 96 well plate in technical triplicates using the ELx808 Absorbance Microplate Reader (BioTek). Precultures of 100 μl were inoculated at an OD600 of 0.1 in microtiter plates and sealed with an air-permeable membrane (Breathe-Easy; Sigma-Aldrich, St. Louis, MO). The growth curves started at OD600 0.1 and incubated at 30°C for 72 h under constant shaking. The OD600 was measured every 10 min.

### Antibodies

The antibody against Sod1 was raised in rabbits using purified recombinant protein. The secondary antibody was ordered from Biorad (Goat Anti-Rabbit IgG (H+L)-HRP Conjugate #172-1019). The horseradish peroxidase-coupled HA antibody was ordered from Roche (Anti-HA-Peroxidase, High Affinity (3F10), #12 013 819 001). Antibodies were diluted in 5% (w/v) nonfat dry milk-TBS (Roth T145.2) with the following dilutions: Anti-Sod1 1:1,000, Anti-HA 1:500, Anti-Rabbit 1:10,000.

### MPP purification

The *E. coli* strain expressing histidine-tagged MPP subunits Mas1 and Mas2 from an expression plasmid was a gift of Vincent Géli (Luciano et al., 1997). The cells were grown at 37°C to an OD600 of 1 and induced with 0.5 mM IPTG over night at 30°C. The bacteria were harvested and resuspended in buffer A (250 mM NaCl, 10 mM imidazole, 2 mM β-mercaptoethanol, 0.2% NP-40, lysozyme), incubated at room temperature for 15 min and snap-frozen in liquid nitrogen. After thawing, DNAseI was added and the cells were sonified 20 times for 1 s at 60% duty level with a Branson sonifier 250. The lysate was cleared by centrifugation, loaded on buffer A-equilibrated Ni-NTA Sepharose resin (Aminitra; Expedeon, San Diego, CA) and washed with buffer A without detergent. A second wash was performed with buffer A adjusted to 1 M NaCl. The enzyme was eluted in buffer C (250 mM NaCl, 300 mM imidazole, 5% glycerol) and stored at −80°C.

### MPP in vitro cleavage assay

The *in vitro* cleavage reactions were performed in 150 mM NaCl, 10% glycerol, 100 μM MnCl2, and 50 mM Tris, pH 7.5, in the presence of 250 μg of purified MPP and 5% reticulocyte lysate containing the substrate. If not otherwise stated, the incubation time was 1.5 h at 30°C. To inhibit MPP, the same reaction was performed in the presence of 2.5 mM EDTA and without addition of MnCl2.

### Prediction of iMTS-Ls

iMTS-Ls were predicted essentially as described (Backes et al., 2018, Boos et al., 2018). Briefly, multiple truncated sequences were generated by sequentially removing amino acids, one by one, N-terminally from the protein of interest. These sequences were submitted to TargetP with appropriate choice of “plant” or “non-plant” organism and without cutoffs. The mTP scores obtained were plotted against the corresponding amino acid position. A Savitzky–Golay filtering step with a window size of 21 (the expected value of the length distribution of known MTSs) was used to smooth the iMTS-L profile. The iMTS-L profiles can also be calculated using the iMTS-L predictor service online tool iMLP (http://imlp.bio.uni-kl.de/).

### Arg5,6 homologs, intracellular localization and genomic distance

Amino acid sequences of yeast Arg5,6 and both post-translationally processed proteins Arg5 and Arg6 were retrieved from (Boonchird et al., 1991a) and used as query sequences for a Diamond BLASTp (Buchfink et al., 2015) against a dataset of 150 eukaryotic genomes (Suppl. Table 1D). Subject organisms that exhibited hits (at least 25% local identity and a maximum E-value of 1E-10) were grouped by the number and pattern of the identified homologs, corresponding to either encoding Arg5,6 as one gene or as two genes (Suppl. Table 1E), by a python script. The intracellular localization of all homologs was checked via TargetP 2.0 (Almagro Armenteros et al., 2019). Results were plotted on a eukaryotic reference tree generated in a previous analysis (Brueckner and Martin, 2020). For the identification of prokaryotic homologs, argB and argC amino acid sequences of *Escherichia coli K-12* were used as queries to search our dataset of 5,655 complete prokaryotic genomes (Refseq, Suppl. Table 1A) (O’Leary et al., 2016). Query sequences were identified as homologs to yeast Arg5 and Arg6 via diamond BLASTp. Only prokaryotic subject organisms exhibiting exactly one homolog to each of the two query sequences (at least 25% local identity and a maximum E-value of 1E-10) were further investigated and the nucleotide distance between the subject genes was calculated (Suppl. Table 1C). Sequence pairs with a maximum distance of 30 nucleotides were suspected to be encoded polycistronically. All genomic distances in nucleotides were derived from Refseq genome feature tables. Sequence pairs with overlapping start and end position were given a distance of one nucleotide.

## Author contributions

F.B. and J.M.H. conceived and supervised the study. J.F. generated constructs and strains and performed *in vivo* experiments. J.F. and E.P. purified recombinant proteins and performed *in vitro* MPP digestions. J.F. and C.G. performed *in vitro* import assays. J.F. and F.B. performed *in silico* prediction of iMTS-Ls. M.R.K. and S.B.G. analyzed the organization of *ARG5,6* homologs in different species. J.F., J.M.H. and F.B. analyzed the data. F.B. wrote the manuscript to which all authors contributed.

## Acknowledgements

We thank Sabine Knaus, Alexander Grevel and Thomas Becker for assistance with the experiments, and Abdussalam Azem and Katja Hansen for helpful discussions and critical reading of the manuscript. This project was funded by grants from the Deutsche Forschungsgemeinschaft (DIP MitoBalance and 2803/10-1 to J.M.H. and 267205415 – SFB 1208 to S.B.G.), the Volkswagen Stiftung (Life) to S.B.G., the Minerva Stiftung (to J.F.) and the Joachim Herz Stiftung (to F.B.).

---

**Supplementary Figure 1.**
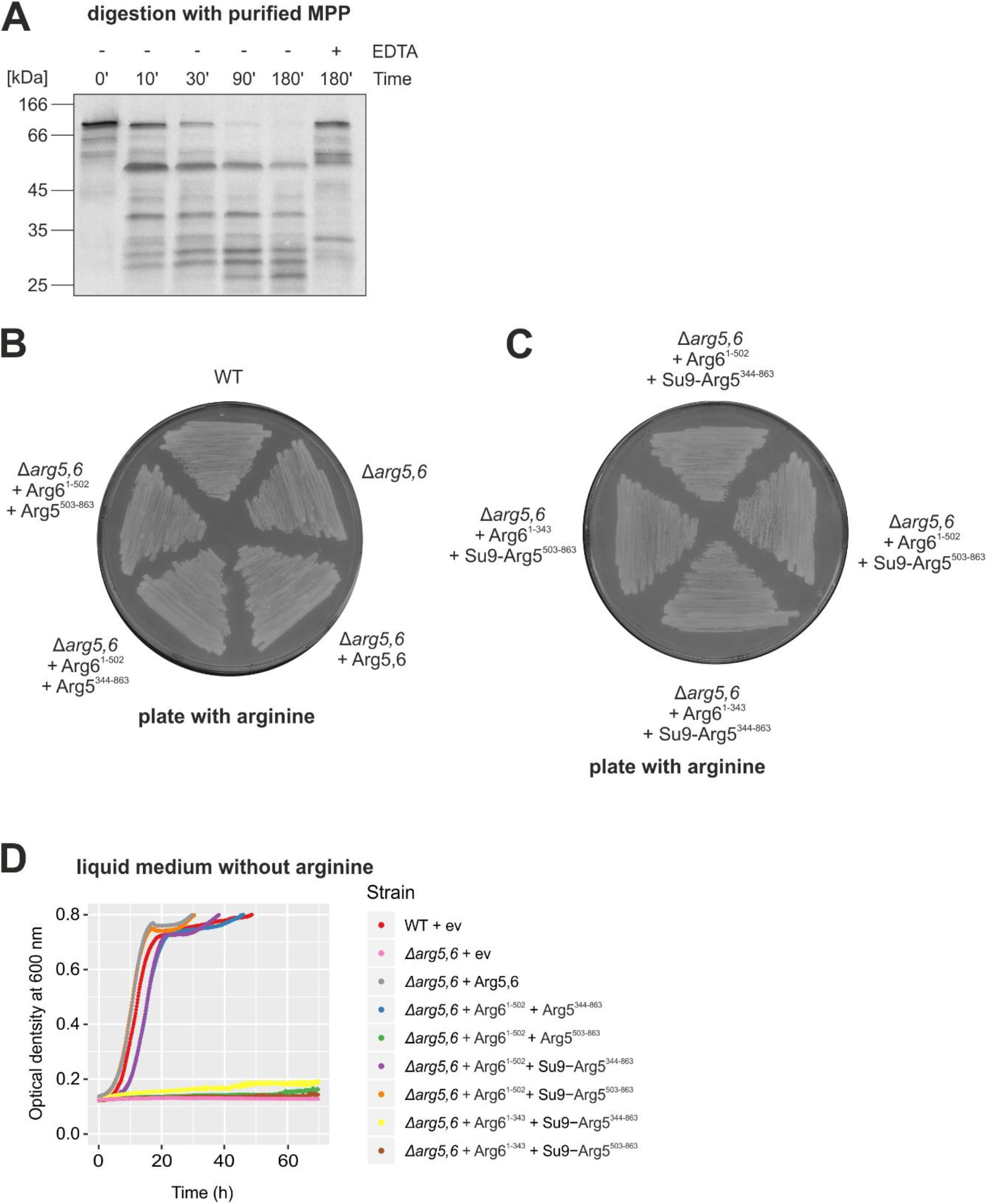
Arg5,6 is internally processed by MPP *in vitro* and separate expression of Arg5 and Arg6 *in vivo* is non-toxic and can restore arginine prototrophy. **A,** Radiolabeled Arg5,6 precursor is incubated with purified MPP for indicated times. EDTA is added as a control to inhibit the enzymatic activity of MPP. Reactions were analyzed by SDS-PAGE, Western blotting and autoradiography. **B-C,** The indicated Arg5,6 variants are expressed in yeast cells lacking *ARG5,6* and cells are streaked out on plates with minimal medium containing arginine. **D,** Cells are grown in liquid minimal medium without arginine for 72 hours at 30°C. The optical density at 600 nm is measured every 10 min.

**Supplementary Figure 2.**
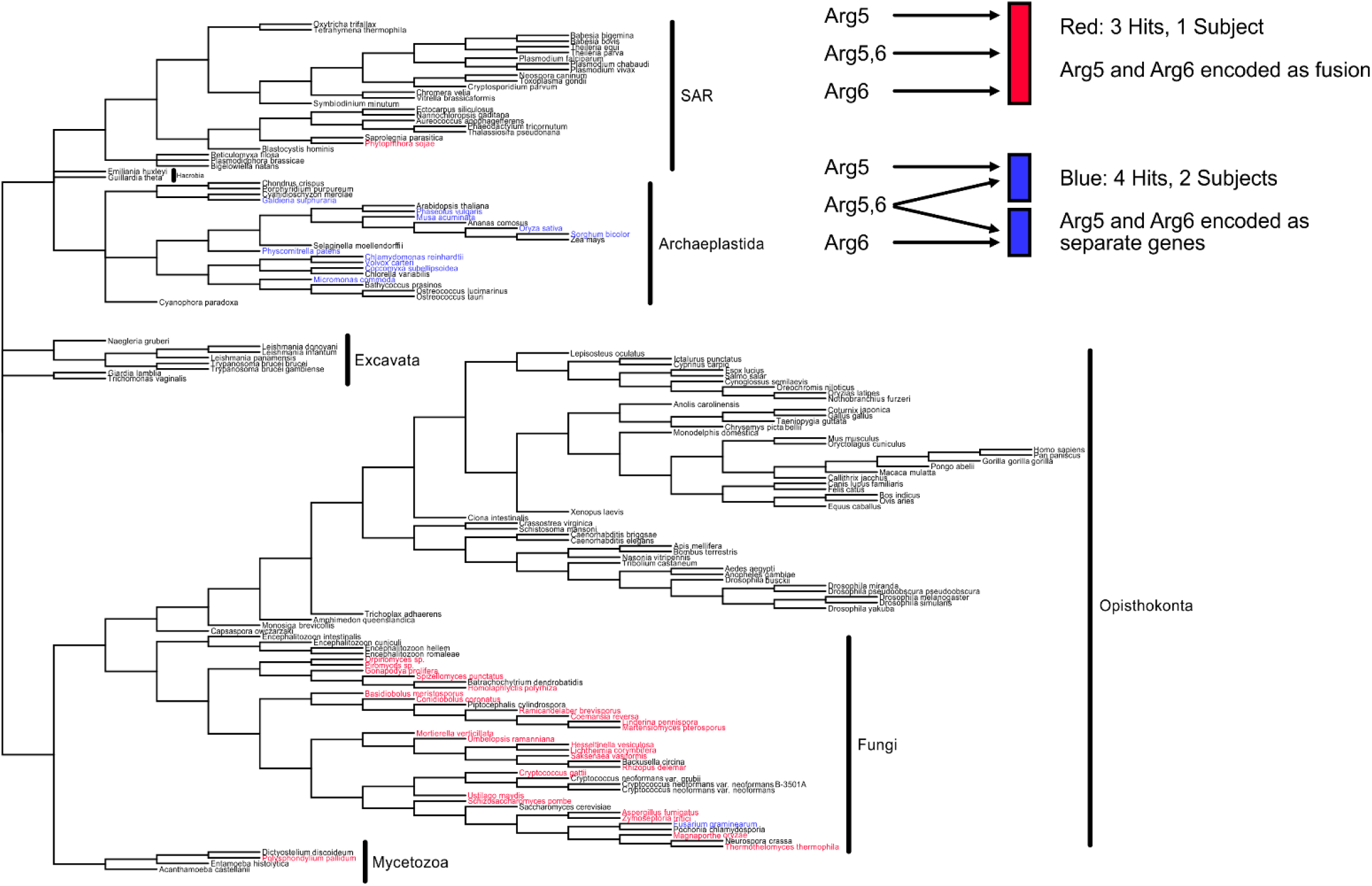
Tandem organization of Arg5,6 is conserved in fungi, whereas algae synthesize two separate proteins. A database of 150 eukaryotes was searched for homologs of full length Arg5,6 and the separate Arg5 and Arg6 proteins of *S. cerevisiae*. Red, species in which the Arg5,6 homolog is encoded as fusion protein. Blue, Arg6 and Arg5 homologs are encoded as separate proteins. Black, no homolog of Arg5,6 found or unclear pattern.

## Supplementary Tables

**Supplementary Table 1.** Evolutionary conservation of the *ARG5,6* gene structure

**Supplementary Table 2.** Yeast strains and plasmids used in this study.

